# A filamentous scaffold for gene regulation

**DOI:** 10.1101/2024.10.07.617013

**Authors:** Tim Rasmussen, Jannik Küspert, Lars Schönemann, Dietmar Geiger, Bettina Böttcher

## Abstract

Proteins of the Drosophila behaviour/human splicing (DBHS) family are involved in many aspects of gene regulation and maintenance like transcription, splicing and DNA repair. The three known members of this family in humans, Non-POU domain-containing octamer-binding protein (NONO), splicing factor proline/glutamine rich (SFPQ), and paraspeckle protein component 1 (PSPC1), form homo- and heterodimers to fulfil these functions by mediating contacts between RNA, DNA, and other protein factors. The dimers can further dynamically oligomerise through α-helical coiled-coils to larger aggregates, which is crucial for many functions of DBHS proteins. While the atomic structures of the dimers are established, the native arrangement in higher oligomers was unknown. Here we present the structure of a filamentous NONO/SFPQ heterooligomer from *Cricetulus griseus* resolved by cryo-EM. Globular heterodimer domains are alternating on both sides of a strand that is stabilized by an interdigitating network of coiled-coil interactions. Two of these strands assemble into a double strand with only few interactions between them. The globular domains of SFPQ face the counter strand and form a groove while those of NONO face outwards. The different environments of NONO and SFPQ in the filament provide the basis for a differential functionality.

**GRAPHICAL ABSTRACT:** 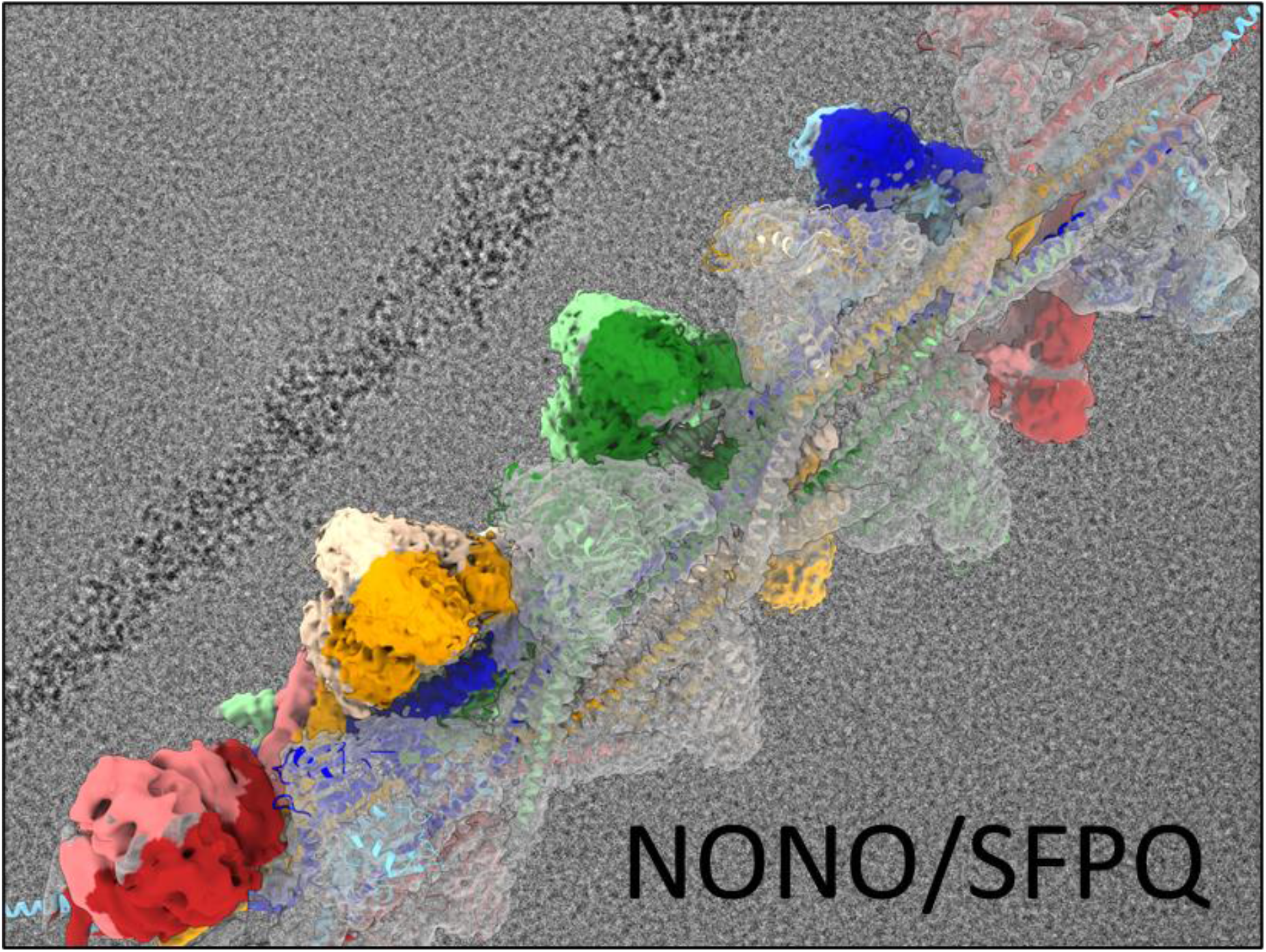

## INTRODUCTION

DBHS proteins are involved in many gene regulation processes at DNA and RNA level (1, 2). These proteins are only found in animals but within this group they are highly conserved. They bind RNA and contribute to RNA-dependent processes. NONO (also abbreviated p54^nrb^) and SFPQ (also called polypyrimidine-tract-binding protein splicing factor, PSF) interact with the C-terminal domain of RNA polymerase II from initiation to termination (3). After transcription, NONO and SFPQ have been implicated in RNA splicing and polyadenylation. For example it has been shown that they are associated with the spliceosome (4–6). DBHS proteins together with long non-coding RNAs like NEAT1 are also seeding and developing subnuclear bodies called paraspeckles (7). They play a role in gene regulation by retaining RNA within the nucleus. In addition, paraspeckles sequester free SFPQ from the nucleoplasm which influences gene expression because SFPQ can bind to DNA and serves as repressor or activator of certain genes (8, 9) which is probably also true for NONO.

DNA double-stranded break repair is another important role that DBHS proteins fulfil. In particular, the nonhomologous end-joining requires NONO which can be substituted by PSPC1 (10) and *in vitro* the purified NONO/SFPQ heterocomplex stimulates end joining (11). Homologous recombination, the other mechanism of DNA double-stranded break repair, involves SFPQ which interacts with RAD51 (12, 13).

Considering the multiple roles of DBHS proteins in gene regulation, it is unsurprising that they are involved in higher functions of the mammalian cell like development, cell cycle and circadian rhythm. Consequently, malfunction leads to severe health problems (14). To just mention two examples, intellectual disabilities can be caused by mutations in NONO (15) due to its role in neuronal cell differentiation and development. In cancer cells NONO is often upregulated (16, 17) and related to a poor survival prognosis (18). In general, many studies found an altered expression of NONO in cancer (19) and also the other DBHS proteins have been connected to cancer so that they may serve as therapeutic target (14, 20).

The molecular organisation of DBHS proteins and their common domains are revealed by sequence alignments and X-ray structures of homo- and heterodimers (21–28). The DBHS core consists of two RNA recognition motifs (RRMs), together with a NonA/paraspeckle domain (NOPS) and a coiled-coil domain (Figure 1A, S1). N- and C-terminal extensions differ between NONO, SFPQ, and PSPC1 and include disordered regions and nuclear localisation signals. In addition, SFPQ possesses an uncharacterised DNA-binding domain N-terminal to the DBHS core. Long α-helices of the coiled-coil domain extend from the globular dimeric domain which inspired models of oligomerisation (21, 29, 30) or showed actual interaction with other dimer units in crystals (31). For SFPQ it was demonstrated that a specific coiled-coil interaction motif in this extension (Figure S1A, region 1) is required for transcriptional regulation, paraspeckle formation and DNA binding (31) highlighting the functional role of oligomerisation.

**Figure 1:**
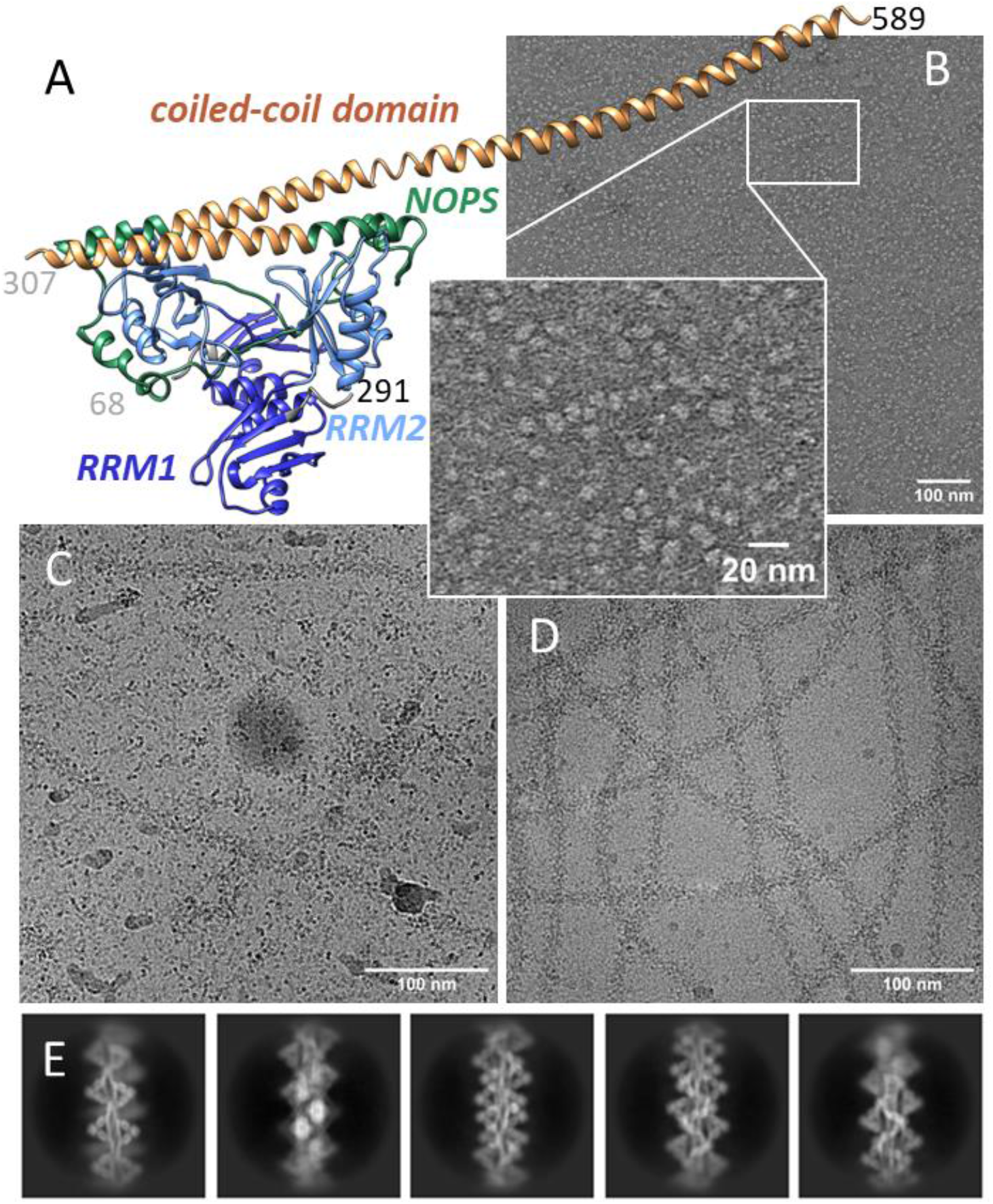
Dimer structure, EM micrographs and 2D classes of NONO/SFPQ. (A) Crystal structure of the NONO/SFPQ heterodimer (PDB: 6WMZ) showing two RNA recognition motives (RRM1, dark blue and RRM2, light blue), a NonA/paraspeckle domain (NOPS, green) and a coiled-coil domain (brown). The numbers of the N-terminal and C-terminal amino acids for NONO (grey) and SFPQ (black) resolved in the structure are marked. (B) Micrographs of negatively stained NONO/SFPQ at a concentration of 50 μg/ml. A few short chains were observed (insert). Micrographs of vitrified samples of NONO/SFPQ at a concentration of 0.2 mg/ml (C) and 1 mg/ml (D). (E) 2D class averages of filaments are shown in (D). The edge of the class images corresponds to 48.4 nm.

The many functional studies on DBHS proteins and interaction studies contrast the limited understanding of the molecular mechanisms behind them. DBHS proteins do not have a catalytic activity on their own. As such their contribution to gene regulation can be understood by providing a common scaffold and binding platform for protein, RNA and DNA. The thorough characterisation of the molecular structure of the DBHS dimers cannot fully address the realisation that the higher oligomerisation of the dimers is crucial to many of their functions. To fill this gap, we present the first polymer DBHS structure consisting of flexible helical filaments.

## MATERIAL AND METHODS

### Purification of the NONO-SFPQ heterodimer

ExpiCHO cells (Gibco) were grown in suspension in Expi293 serum-free medium (Gibco). For expression, a 1 l (4x 250 ml) suspension culture was inoculated to 0.75 x106 cells/ml. The culture was harvested after 96 h, yielding a cell pellet of about 9 g. The cell pellet was frozen and stored at -20°C until further use. For purification, the cells were suspended and lysed in a Teflon-in-glass homogeniser in 50 ml of lysis buffer containing 1 % dodecylmaltoside (DDM, Glycon Biochemicals, Germany), 50 mM sodium phosphate buffer pH 7.5, 300 mM NaCl, 10 % glycerol, 20 mM imidazole, complete EDTA-free protease inhibitor cocktail (Roche) and 1 mM phenylmethanesulfonyl fluoride at 4°C. After 1 hour of incubation on ice, 4 mg DNAse (Roche) and 4 mM MgCl_2_ was added and incubated for another hour. After 60 min centrifugation at 30,000 g, the supernatant was applied to a 1 ml Ni nitrilotriacetic acid (Ni-NTA) agarose column (Sigma) beforehand equilibrated with washing buffer, containing 0.05 % DDM, 50 mM sodium phosphate buffer pH 7.5, 300 mM NaCl, 10 % glycerol and 20 mM imidazole. A native histidine-rich motif of NONO (HHQHHHQQHH, residues 19 to 28 in CgNONO) is employed as native binding motif for this purification step. The column was washed with 40 ml washing buffer and proteins eluted in 1 ml fraction with an elution buffer, composed the same as the washing buffer but with 300 mM imidazole. The peak fraction was applied to a 10/300 Superdex200 Increase column (Cytiva) equilibrated with SEC buffer, containing 50 mM HEPES pH 7.5, 150 mM NaCl, 5 mM EDTA and 0.03 % DDM. Samples were analysed by SDS-PAGE using hand cast 15 % acrylamide gels. The two main bands on a Coomassie stained gel were cut and identified by mass spectrometry in the Rudolf-Virchow-Zentrum MS facility (University Würzburg) after tryptic digest as SFPQ and NONO, respectively. A Western blot on nitrocellulose using a PentaHis Antibody HRP conjugate (Qiagen) produced a positive result for the band identified as NONO. The main peak from the size exclusion chromatography was further concentrated with Vivaspin 100 kDa cutoff concentrators (Millipore) and 10 mM MgCl_2_ was added before vitrification.

### Electron microscopy and data analysis

#### Negative stained samples

Copper grids with a continuous carbon film were glow discharged for 150 sec before use. The sample was diluted to 50 μg/ml with SEC buffer and incubated on the grid for 1 min. After washing with water, the grids were washed twice with 2 % uranyl acetate followed by incubation with 2% uranyl acetate for 5 min and removal of excess liquid. Stained samples were imaged in a FEI Tecnai T12 electron microscope at a nominal 52,000x magnification, with an exposure of 30 electrons/Å^2^ and a target defocus of 1 μm.

#### Vitrified samples

Samples were concentrated to 0.2 mg/ml or 1 mg/ml and prepared either on copper grids with holey carbon support and an additional layer of 2 nm continuous carbon (R1.2,1.3, Quantifoil) or on UltAufoil grids (R0.6/1.0, Quantifoil), respectively. The grids were glow discharged for 60 sec or 150 sec, respectively, in air at a pressure of 3.0×10^−1^ Torr at medium power with a Harrick Plasma Cleaner (PDC-002) and used within one hour for vitrification. The samples were vitrified in a Vitrobot IV in ethane with 5 s blot time and +20 blot force. Movie data in EER format was acquired on a Krios G3 electron microscope (ThermoFisher) equipped with a Falcon IVi direct detector and a Selectris energy filter (cryo-EM facility Würzburg). The zero-loss movies were recorded with a slit width of 5 eV at a magnification of 130,000x with a calibrated pixel size of 0.946 Å and a total exposure of 70 electrons/Å^2^ (exposure time of 6.2 s). The target defocus range was between 0.5 to 1.4 μm for the concentrated sample and between 1.4-3 μm for the dilute sample on the grids with an additional 2 nm carbon film. For single particle image processing, 17059 movies of the concentrated sample were recorded in EER format and motion corrected and dose weighted in 40 fraction in a live session of the program package *CryoSparc* version 4.5 (32). All subsequent steps of image processing were also performed in *CryoSparc*, starting with the patch based CTF estimation. 4.2 million filament segments were picked with the blob picker. The segment images were further cleaned in two rounds of 2D classification followed by the selection of the best classes, retaining 3 million segment images. The initial models were determined by helical refinement and gave a resolution of 5.9 Å. The asymmetric non-uniform refinement of the same segments provided a map with an overall resolution of 3.9 Å. A 3D classification in *CryoSparc* was performed into 10 classes but further refinement of subpopulations did not improve resolution. Local refinements focused on 8 or 4 dimers on one strand excluding the RRM1 domains and resulted in a resolution of 3.5 or 3.3 Å, respectively. The required masks were obtained by fitting a crystal structure into the map (see below) and using the “colour zone” tool in program *Chimera* version 1.17.3 (34) to select specific regions. Both strands were refined separately and combined with the overall map to a consensus map using the combine focused maps tool of the program *Phenix* version 1.20.1 (33). The crystal structure of the human NONO/SFPQ heterodimer (PDB: 6WMZ) was fitted rigidly into the dimer locations in the filament using the program *Chimera*. The model was manually optimised using *Coot* version 0.9.8.93 (35, 36) in the locally-refined maps with 4 subunits for residues 148 to 340 of NONO and 367 to 594 of SFPQ which excludes the RRM1 domains. The models were refined with the realspace refinement tool of *Phenix*. An overall model of the composite map included the RRM1 domains, which were fitted as rigid bodies. Complete dimer models from the local refinement were used as templates to obtain four repeats within the filament by rigid body fitting to the composite map, visualising the overall biological assembly. Finally, the model was refined with *Phenix* realspace refinement. Map and model parameters are summarised in Table S1. Figures were prepared and subunits compared (matchmaker) in *ChimeraX* (37). Sequence alignments were computed with *M-coffee* (38). The isoforms X1 were taken for sequence annotation of NONO (NCBI Reference Sequence: XP_007649790.2) and SFPQ (NCBI Reference Sequence: XP_035310744.1).

## RESULTS

### Dimers of NONO/SFPQ heterodimers form concentration dependent filaments

NONO and SFPQ are highly conserved between humans and Chinese Hamster (*Cricetulus griseus)* with an identity of 98% (Figure S2,S3). The native full-length NONO/SFPQ was purified from Chinese Hamster Ovary Cells (*Cricetulus griseus*; Figure S1B,C) and the purified protein was verified by tryptic digest and mass spectrometry. Electron micrographs of negatively stained NONO/SFPQ at a concentration of 50 μg/ml showed regular particles with a diameter of about 10 nm and some short chains of several of these particles (Figure 1B). Micrographs of vitrified samples at a higher concentration of 0.2 mg/ml showed similar particles but also some longer filaments (Figure 1C). Eventually, at a concentration 1 mg/ml the individual particles disappeared and only filaments were observed (Figure 1D).

### NONO/SFPQ filaments consist of a flexible double-stranded helix

The filaments were inherently flexible with a varying local curvature (Figure 1D). All filaments had the same diameter of about 15 nm, which is also evident from 2D-class averages (Figure 1E). The filaments of the NONO/SFPQ-dimers consisted of two intertwined strands with globular domains alternating on both sides of each strand (Figure 2). Within one of the intertwined strands the heterodimers on opposing sides were arranged antiparallel and both strands of the filament are also antiparallel. The filaments had a left-handed twist of about -20° and a rise of 10 nm with an asymmetric unit of two heterodimers per strand (Figure 2). That corresponds to a helical pitch of around 180 nm with a total of 36 NONO/SFPQ dimers per turn.

**Figure 2:**
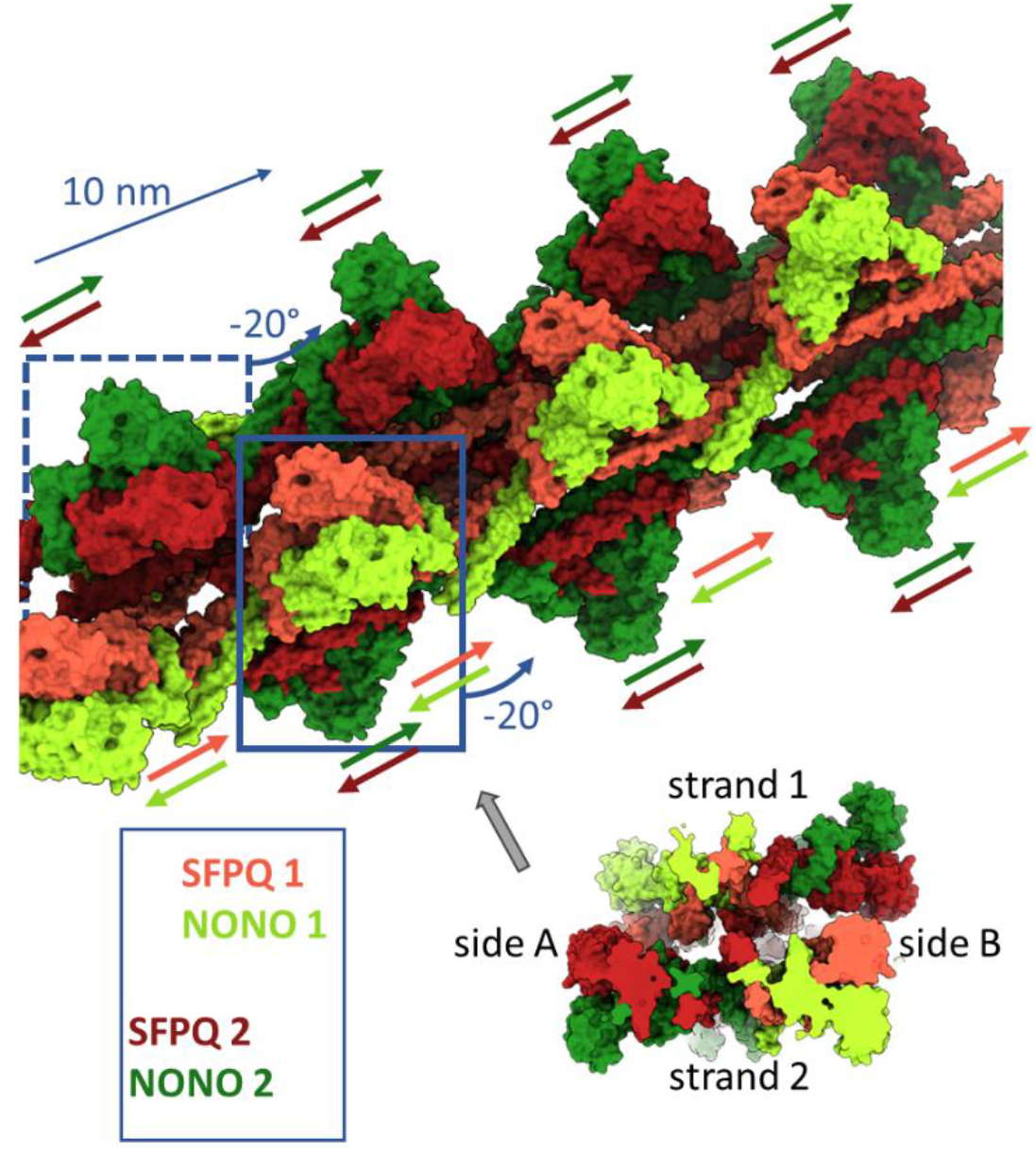
Filamentous structure of the NONO/SFPQ heterocomplex. (A) NONO/SFPQ form a left-handed helical double strand with a turn of -20° and a rise of 10 nm. SFPQ (red) is always facing the other strand while NONO (green) is facing outwards. The filament structure can be built up from two non-equivalent dimer blocks on each strand (blue squares). The direction of the long α-helices from N-to C-terminus is indicated by the coloured arrows, showing that the non-equivalent dimers are orientated in opposite directions along the filament. The insert shows a view along the filament (position grey arrow).

Overall, a resolution of 3.9 Å was obtained with large local variations. The protruding globular domains were less well resolved than the interior of the filament (Figure S4). These globular domains had a similar tetrahedral shape as the RRM containing domains of NONO/SFPQ in the crystal structures (27, 39)(Figure 1A). The inherently flexible RRM1 at the tips of the globular domains coincided with the regions of lowest local resolution in the map (Figure S4).

Considering a possible movement within each strand and against each other as well as the probable inherent flexibility of the RRM1 domains, we did local refinements of 8 or 4 heterodimers from each strand excluding the RRM1 domains. This provided maps for the individual strands with resolutions of 3.5 or 3.3 Å, respectively (Table S1). The higher resolution refinements with the central 4 subunits were modelled excluding the RRM1 domains (Figure S5). All local refinements were combined to a composite map of the double strand. The map of the double strand accounted for residues 68-340 of NONO and 287-593 of SFPQ, including the rigidly fitted RRM1 domains, while the N-terminal and C-terminal low-complexity regions were not resolved.

The flexibility was further characterised with a 3D classification which confirmed bending of the filament, predominantly visible when looking in the direction where one strand is in front and one in the back (Figure S6; Movie 1).

### Coiled-coils provide a strong network within a strand

The long α-helical regions beyond the globular domains had the highest local resolution and allowed confident modelling of the interactions between the different heterodimers. Within one strand, we identified three regions of interaction (region 1-3) which were mediated by coiled coils (Figure 3).

**Figure 3:**
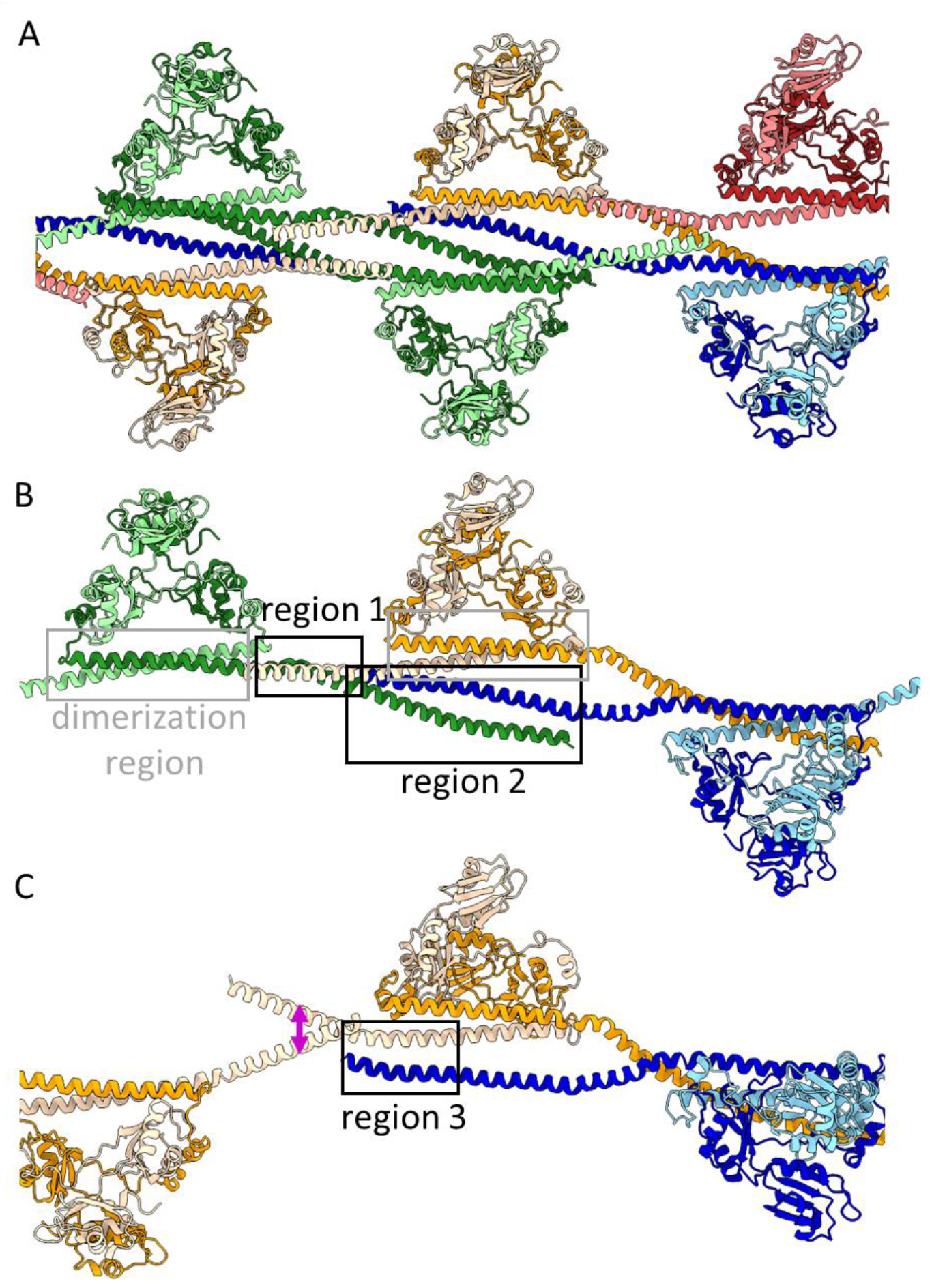
Interactions between dimer building blocks within one strand. (A) The model of one of the two strands is shown for highlighting the α-helical network. The heterodimers have different colours with NONO in light and SFPQ in darker colours. (B) The coiled-coil domain engages in interactions within a dimer (dimerization region, grey box) and with other dimers within a strand. Region 1 provides interaction with the direct neighbour on the same side of the strand while region 2 interacts with a dimer further away on the opposite side of the strand. (C) There is a crossover of NONO α helices (pink arrow) and in region 3 another short helical coiled-coil which provide additional interactions between dimers different to the ones shown in (B).

In region 1, close to the globular RRMs, NONO and SFPQ formed a left-handed antiparallel coiled-coil that was stabilized by a leucin zipper. A coiled-coil has also been described for SFPQ homodimers in crystal structures for this region (31) but in our filaments the kink of the α-helix of SFPQ between the globular domain and region 1 is more pronounced (Figure S7A). Region 1 corresponds to residue 524-551 in SFPQ and to residue 303-330 in NONO. Over the total length of 4 nm, both helices form multiple hydrophobic interactions and salt bridges. Of notice, at one end close to the kink of SFPQ, H524 (SFPQ) forms a π-π-interaction with H330 of NONO. In the middle of region 1 R538 of SFPQ and R319 of NONO interact (Figure 4A). Despite the likewise charge, such arginine-arginine contacts frequently promote protein-protein interactions and are energetical favourable (40, 41). An additional arginine-arginine bridge is established between R287 of NONO and R568 of SFPQ away from region 1.

**Figure 4:**
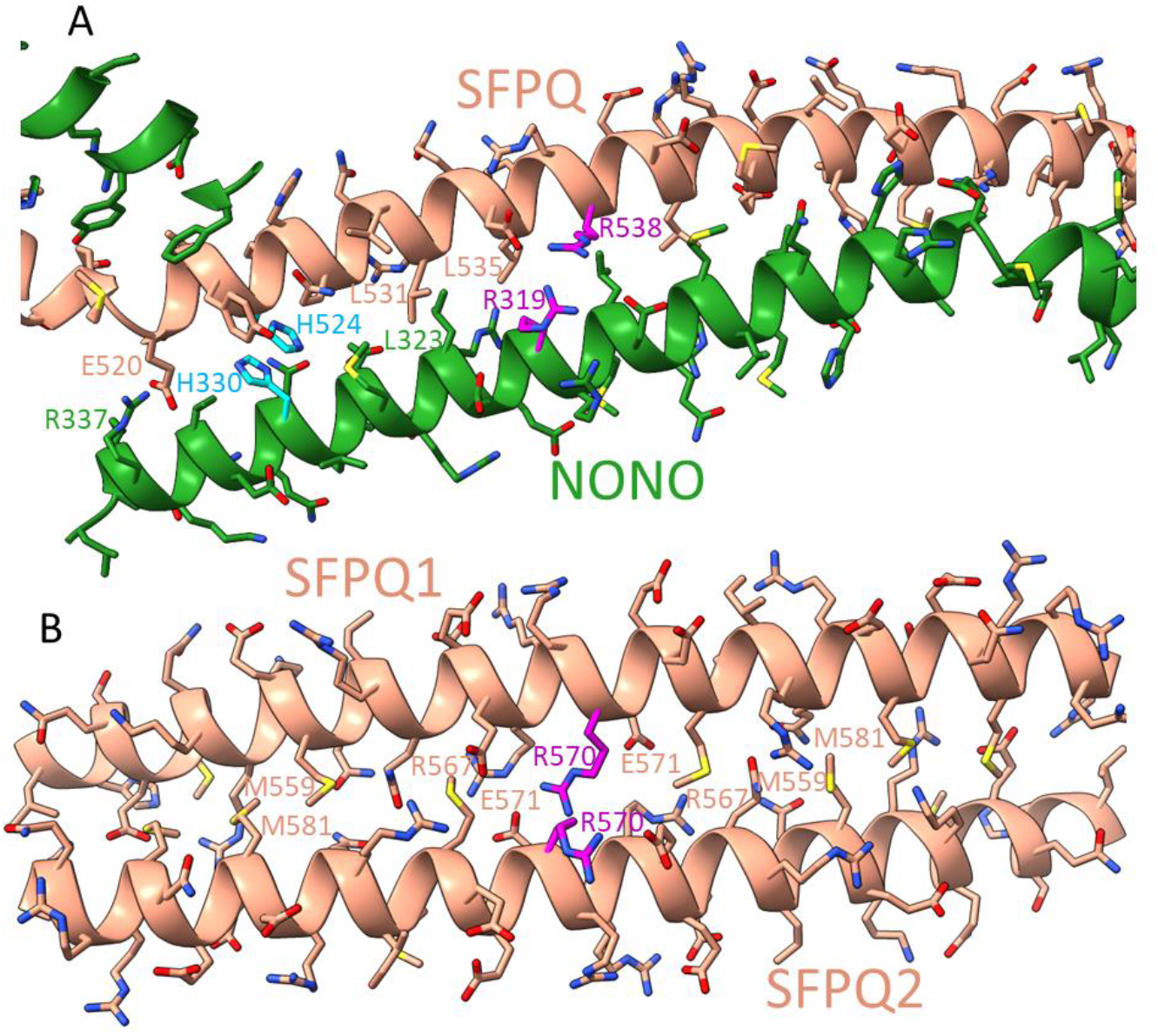
Coiled-coil interactions. (A) Interaction region1: The long α-helices of SFPQ and NONO from neighbouring dimers establish a coiled-coil. Some residues important for this interaction are marked. For example, R319 (NONO) interacts with R538 (SFPQ) in an arginine-arginine bridge (purple) and H330 (NONO) and H524 (SFPQ) engage in a π-π stacking interaction (cyan). (B) Interaction region 2: The long α-helices of non-equivalent SFPQ on opposite sides of the same strand form a coiled-coil. Where opposing R570 residues meet, the SFPQ subunits have different conformations, else the side chains would clash (purple).

At the end of region 1 close to residue 550 in SFPQ, starts region 2. It promotes the interaction with another SFPQ subunit on the opposite side of the same strand but two heterodimers away (Figure 3B). Over a distance of 6 nm both SFPQ subunits form a right-handed antiparallel coiled-coil from residue 548-589, which is stabilized by intercalating methionine residues. Like in region 1, the centre of region 2 is formed by a cluster of charged residues (R567, R570, and E571) from the opposing SFPQs. It is notable that the side chain conformations of the two R570 differ (Figure 4B). Taken together, the long α-helix of SFPQ first interacts with NONO of the same heterodimer (dimerization domain) in a right-handed coiled-coil, then with NONO of the neighbouring heterodimer in a left-handed coiled-coil (region 1; Figure 4A), and finally with another SFPQ in a right-handed coiled-coil (region 2).

An additional short interaction site (region 3, Figure 3C) overlaps partly with region 2 in SFPQ and mediates the contact to NONO between the dimerization domain and region 1 in the opposing heterodimer: Here, the closest contacts are R589 (SFPQ) with H305 (NONO) and M575 (SFPQ) with M298 (NONO). Region 3 runs from R568 to R589 in SFPQ and from D283 to H305 in NONO and coincides with the kink on the long α-helix in NONO at E299. An additional crossover of the α-helices from NONO of different heterodimers is stabilized by a salt bridge between R324 and E327/E328 (Figure 3C).

### Weak interactions between strands and ordered subunit arrangement

It is well established that NONO and SFPQ preferentially form heterodimers (23, 27, 28, 31). The isolated dimer has a pseudo twofold symmetry with equivalent positions of SFPQ and NONO. In the context of the double stranded filaments, the SFPQ faces the other strand while NONO is on the periphery (Figure 2). Because of the ordered arrangement of the subunits, only SFPQ and not NONO contributes to the weak interactions between the strands. The EM density suggests that the end of the long α-helix of SFPQ interacts with the NOPS domain at the loop Q461 to P464 of the opposing strand (Figure S8). It is notable that this loop deviates from NONO in 4 residues and represents the largest sequence difference between SFPQ and NONO in the common core region (Figure S9). Another relevant difference is an insertion of 4 residues in region 2 in SFPQ, which extends the coiled-coil helix by about 0.6 nm compared to NONO and brings it into contact with the NOPS domain. Together, both sequence differences are sufficient to promote the contact between two SFPQs but not between two NONOs or a NONO and a SFPQ on opposing strands. However, this contact is present only in half of the heterodimers on one side of the strand. For the other half, the contact to NOPS is blocked by the same α-helix of the other strand (Figure S10). Despite the differences of the interstrand contact very little difference can be seen for the non-equivalent SFPQ subunits except the above-mentioned side chain clash of R570. An overlay of all SFPQ subunits showed no significant difference in the backbone (RMSD values below 0.53 Å; Figure S11) which was also the case for NONO (RMSD values below 0.44 Å).

## DISCUSSION

Based on the first crystal structures of dimers (21, 31) it was suggested that DBHS proteins form molecular scaffolds for many gene regulation processes (1). A dynamic equilibrium between dimers and higher oligomers depends on the contacts between the long α-helices (31). Here, we describe how these coiled-coils and additional contacts between NONO/SFPQ heterodimers lead to the concentration dependent formation of double-stranded filaments.

While the globular dimer region of the crystal structures fit well into the corresponding densities of the filaments, the kink of the long SFPQ α-helix between the dimerization region and region 1 at residue S517 is much more pronounced in the filaments than in the crystal structures (PDB: 4WIJ, 4WIK, and 6WMZ; only these crystal structures have α-helices beyond the globular domain) (Figure S7A). After the kink the helix continuously bends. NONO shows a similar kink at E299 but the helix is not bending. This part of the α-helix of NONO has so far not been resolved in the crystal structures. The kinks in the long coiled-coil helices of NONO and SFPQ lead to different arrangements of the heterodimers in the filament compared to the packing in the crystal structures. Interesting is the comparison to the crystal structure of a SFPQ homodimer (PDB: 4WIJ) where two dimers have the same distance and offset to each other like two dimers on opposite sides of the strand in the filament (Figure S7B). This is a surprising similarity as there are no direct contacts between these two dimers. In addition, a similar distance to another neighbouring dimer is established through the coiled-coil in region 1. Aligning the coiled-coils from this crystal structure and our filament structure on SFPQ (residues 523-545), reveals that the helices in the filament are closer to each other and have a more pronounced coiling than in the crystal (Figure S7C). This suggests that the interaction in region 1 between NONO and SFPQ is energetically more favourable than in the SFPQ homooligomer and promotes a heteromerization. Such a thermodynamical preference for heteromers is one prerequisite for the ordered overall structure of the filament with NONO on the outside and SFPQ at the inside. The intimate interaction may also promote the kinking of the α-helices at both ends of region 1 where the residues are incompatible with continuing the coiled-coil and are better suited to form other pairs of coiled-coils in the dimerization domain and in region 2.

Recently the structure of the human SFPQ homodimer in complex with RNA showed a considerable conformational change upon RNA binding, especially in the RRM1 region (26). A similar structural rearrangement and flexibility was also indicated by SAXS measurements of the NONO homodimer interacting with RNA (28). Aligning the structure of the SFPQ-RNA complex (26)(PDB: 7UJ1) with the globular region in our filaments shows that the RNA binding site of SFPQ is close to the interstrand contact within the groove (Figure 5A,B) and shielded by the globular domains of RRM1. The second RNA binding site in the dimer is asymmetric to the first in the crystal structure of the SFPQ homodimer (26) and would face the outside of the filament. These outward facing binding sites of NONO form four continuous bands of positively charged grooves that are accessible at the outside of the filaments (Figure 5B,C).

**Figure 5:**
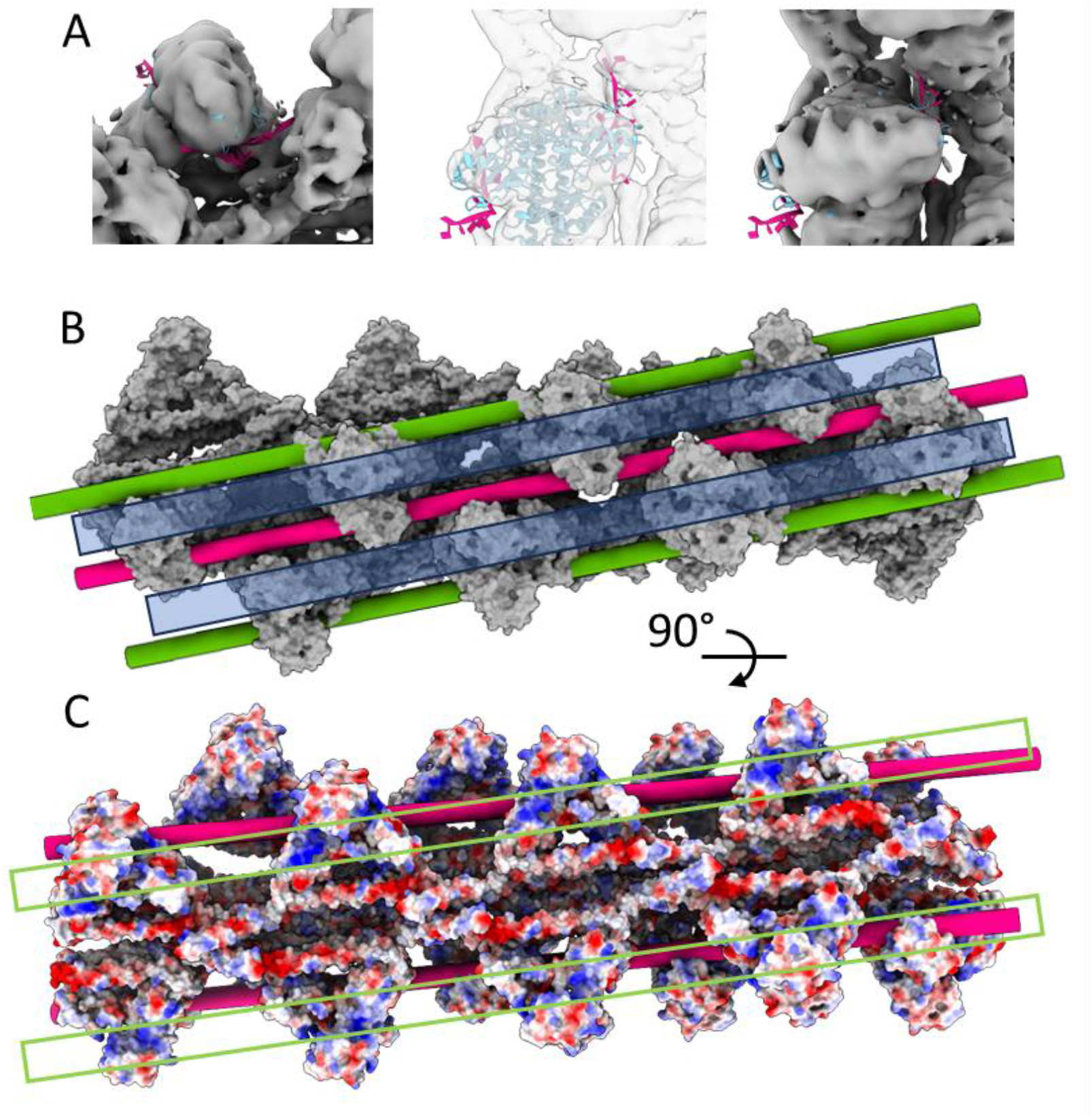
Possible interactions of RNA and DNA with the filaments. (A) The crystal structure of the SFPQ homodimer shows the interaction with RNA between RRM1 and 2 on each subunit (PDB 7UJ1) (Wang et al., 2022). This crystal structure (shown as model) was rigidly fitted into the globular domain of the EM filament map. This would place the RNA (pink) bound to one subunit into the groove between the strands. The other bound RNA on the other subunit faces outwards. An image with solid and with transparent EM density is shown and a different view. (B) Surface representation of NONO/SFPQ model (grey). RNA (pink) could be locked in between the two strands interacting with the binding sites of SFPQ which have opposite directions on both strands. Alternatively, RNA could bind on the periphery to the binding sites of NONO (green). The unresolved DNA binding domain of SFPQ is N-terminal to RRM1 and thus probably close to the top of the heads (blue box). (C) Electrostatic surface representation of the model viewed onto one strand (about 90° to view in (B)). Positively charged grooves of NONO continuing from one head to the next is the likely location for RNA binding (green boxes).

A structure in complex with DNA has not been published yet nor the DNA binding domain has been resolved. This domain is N-terminal to the RRM1 domain on SFPQ and thus likely to be located on top of the globular domains suggesting that DNA could run along the globular heads on one strand (Figure 5B). However, dsDNA is rigid and can only contact two or three consecutive heads in the filament before it diverges from the filament as its persistence length of 40-60 nm does not allow it to follow the twist of the filament. This could explain how the double-stranded NONO/SFPQ helix holds two ends of DNA for double-strand break repair (42), while longer continuous stretches of dsDNA cannot bind.

In summary, we show how NONO/SFPQ heterodimers form double-stranded helical filaments in the absence of RNA or DNA. These filaments provide a binding platform of high avidity which supports the multiple functions of DBHS proteins as a scaffold. Despite the high identity between NONO and SFPQ in the core they have different environments within the double stranded filaments because SFPQ faces the other strand while NONO is fully exposed to the outside.

## Supporting information

Supplemental Tabel and Figures

## DATA AVAILABILITY

The atomic model for the local refinements of the central units has been deposited in the Protein Data Bank under the accession code 9GLC and 9GLD. A model of a composite map visualising better the biological assembly has the accession code 9GNI. The EM map of the overall asymmetric non-uniform refinement (EMD-51413), local refinements of the strands (EMD-51417 and 51418), local refinements of the central units on each strand (EMD-51438 and 51439) and the composite map (EMD-51471) have been deposited in the Electron Microscopy Data Bank (EMDB).

## ACKNOWLEDGEMENTS

We thank Christian Kraft and Michelle Endres for technical assistance. We thank Stephanie Lamer and Andreas Schlosser from the mass spectrometry facility at the Rudolf-Virchow-Zentrum. Electron Cryo Microscopy was carried out in the cryo EM-facility of the Julius-Maximilians-Universität Würzburg.

