## Supplemental Tabel and Figures for "A filamentous scaffold for gene regulation"

**Table S1: Cryo-EM data collection, refinement and validation statistics.**

| | overall | Focused strand 1 | Focused strand 2 | Focused central strand 1 | Focused central strand 2 | Composite Structure |
|---|---|---|---|---|---|---|
| EMDB ID | EMD-51413 | EMD-51417 | EMD-51418 | EMD-51438 | EMD-51439 | EMD-51471 |
| PDB ID | | | | 9GLC | 9GLD | 9GNI |
| **Data collection and processing** | | | | | | |
| Magnification | | | 130,000 | | | |
| Voltage (kV) | | | 300 | | | |
| Electron exposure (e–/Å$^2$) | | | 70 | | | |
| Defocus range (μm) | | | 0.5 to 1.4 | | | |
| Pixel size (Å) | | | 0.946 | | | |
| Symmetry imposed | | | C1 | | | |
| Initial particle images (no.) | | | 4,144,650 | | | |
| Final particle images (no.) | | | 2,974,535 | | | |
| Map resolution (Å) | 3.9 | 3.5 | 3.5 | 3.3 | 3.3 | -- |
| FSC threshold | | | 0.143 | | | |
| **Refinement** | | | | | | |
| Initial model used (PDB code) | -- | -- | -- | 6WMZ | 6WMZ | 6WMZ, 9GLC, 9GLD |
| Model resolution (Å) | -- | -- | -- | 3.7 | 3.8 | 5.9 |
| FSC threshold | | | | 0.5 | 0.5 | 0.5 |
| Model composition | | | | | | |
| Non-hydrogen atoms | | | | 14,742 | 14,629 | 74,499 |
| Protein residues | | | | 1764 | 1751 | 9,027 |
| Ligands | | | | 0 | 0 | 0 |
| *B* factors (Å$^2$) | | | | | | |
| Protein | | | | 109 | 105 | 150 |
| R.m.s. deviations | | | | | | |
| Bond lengths (Å) | | | | 0.005 | 0.007 | 0.005 |
| Bond angles (°) | | | | 0.807 | 0.904 | 1.047 |
| Validation | | | | | | |
| MolProbity score | | | | 2.18 | 2.09 | 1.82 |
| Clashscore | | | | 14.79 | 15.87 | 15.68 |
| Poor rotamers (%) | | | | 3.62 | 2.52 | 0.95 |
| Ramachandran plot | | | | | | |
| Favored (%) | | | | 97.65 | 97.63 | 97.38 |
| Allowed (%) | | | | 2.35 | 2.37 | 2.62 |
| Disallowed (%) | | | | 0.00 | 0.00 | 0.00 |

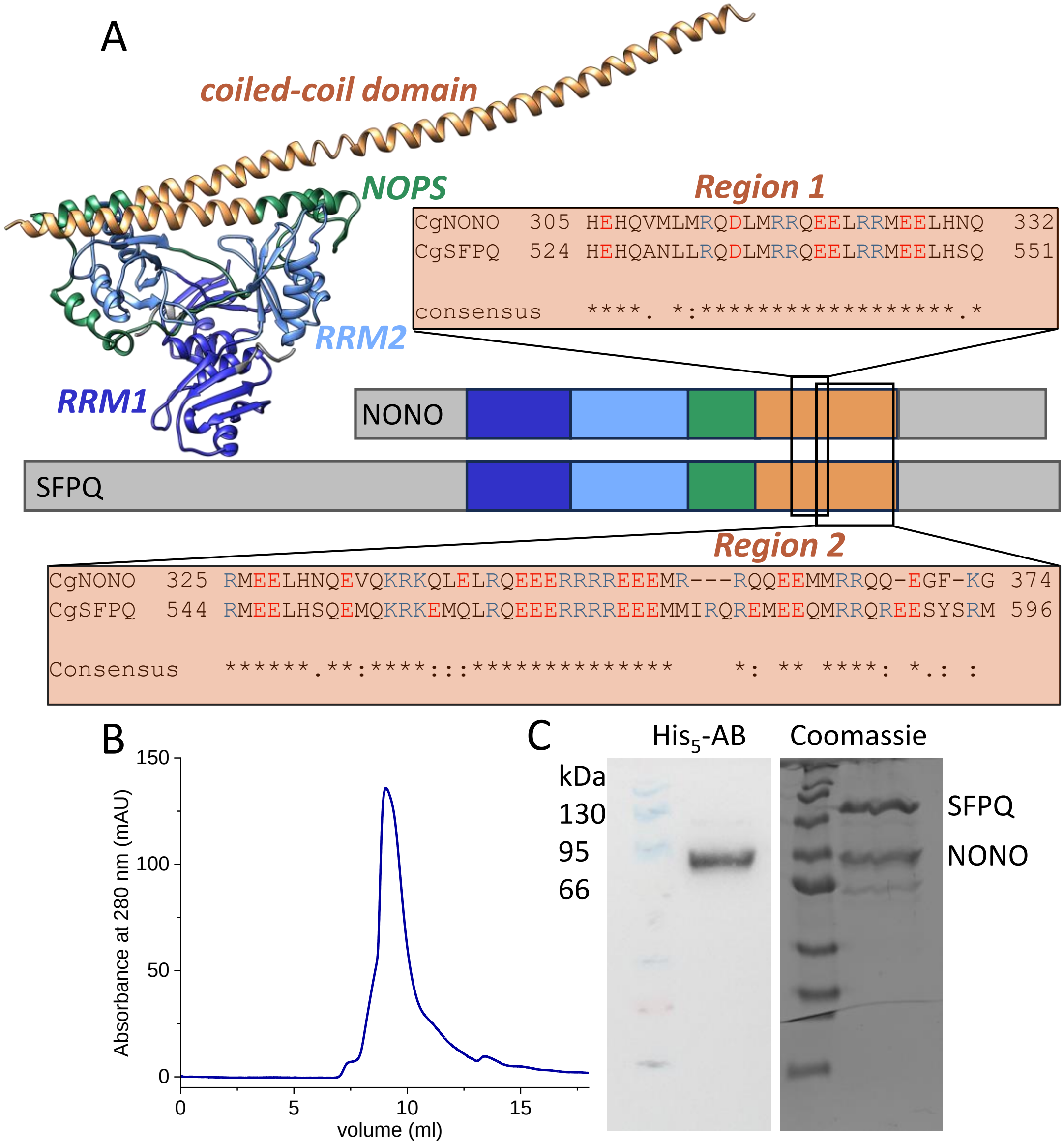


**Figure S1: NONO-SFPQ dimer organisation and purification.** (A) The DBHS core is organised in several common subdomains as shown schematically for the primary sequences for NONO and SFPQ and the crystal structure of the heterodimer (PDB: 6WMZ). Each subunit contributes two RNA recognition motives (RRM1, dark blue and RRM2, light blue), a NonA/paraspeckle domain (NOPS, green) and a coiled-coil domain (brown). The subdomains form a tetrahedral globular domain and the extended α-helices of the coiled-coil domain interact with other dimeric building blocks to oligomerise to a strand. These interactions are provided by region 1 with the neighbouring dimer and by region 2 with a dimer further away. Only SFPQ has dimer-dimer interactions via region 2. While the sequence of region 1 is almost identical between NONO and SFPQ, there is a substantial difference at the C-terminal end of region 2. (B) Size exclusion chromatography of shows an elution for NONO-SFPQ in a distorted main peak. This and the size calibration suggests that not isolated dimers are predominately present at this stage but also not polymeric filaments. (C) The SDS-PAGE shows two main bands in the Coomassie-stained gel at about 80 and 110 kDa which were identified by mass spectrometry as NONO and SFPQ, respectively (right). The former showed a positive detection with a His5-antibody on Western blots (left).

```
hamster_NONO       1 MQSNKTFNLEKQNHTPRKHHQHHHQQHHQQQQQQQQPPPPIPANGQQASSQNEGLTIDLKNFRKPG   66
human_NONO         1 MQSNKTFNLEKQNHTPRKHHQHHHQQQHHQQQQQQPPPPPIPANGQQASSQNEGLTIDLKNFRKPG   66
cons               1 **************************:*:****** ******************************   66
                                                      RRM1
hamster_NONO      67 EKTFTQRSRLFVGNLPPDITEEEMRKLFEKYGKAGEVFIHKDKGFGFIRLETRTLAEIAKVELDNM  132
human_NONO        67 EKTFTQRSRLFVGNLPPDITEEEMRKLFEKYGKAGEVFIHKDKGFGFIRLETRTLAEIAKVELDNM  132
cons              67 ******************************************************************  132
                                                      RRM2
hamster_NONO     133 PLRGKQLRVRFACHSASLTVRNLPQYVSNELLEEAFSVFGQVERAVVIVDDRGRPSGKGIVEFSGK  198
human_NONO       133 PLRGKQLRVRFACHSASLTVRNLPQYVSNELLEEAFSVFGQVERAVVIVDDRGRPSGKGIVEFSGK  198
cons             133 ******************************************************************  198
                                                                 NOPS
hamster_NONO     199 PAARKALDRCSEGSFLLTTFPRPVTVEPMDQLDDEEGLPEKLVIKNQQFHKEREQPPRFAQPGSFE  264
human_NONO       199 PAARKALDRCSEGSFLLTTFPRPVTVEPMDQLDDEEGLPEKLVIKNQQFHKEREQPPRFAQPGSFE  264
cons             199 ******************************************************************  264
                                              coiled-coil domain
hamster_NONO     265 YEYAMRWKALIEMEKQQQDQVDRNIKEAREKLEMEMEAARHEHQVMLMRQDLMRRQEELRRMEELH  330
human_NONO       265 YEYAMRWKALIEMEKQQQDQVDRNIKEAREKLEMEMEAARHEHQVMLMRQDLMRRQEELRRMEELH  330
cons             265 ******************************************************************  330

hamster_NONO     331 NQEVQKRKQLELRQEEERRRREEEMRRQQEEMMRRQQEGFKGTFPDAREQEIRMGQMAMGGAMGIN  396
human_NONO       331 NQEVQKRKQLELRQEEERRRREEEMRRQQEEMMRRQQEGFKGTFPDAREQEIRMGQMAMGGAMGIN  396
cons             331 ******************************************************************  396

hamster_NONO     397 NRGAMPPAPVPTGTPAPPGPATMMPDGTLGLTPPTTERFGQAATMEGIGAIGGTPPAFNRPAPGAD  462
human_NONO       397 NRGAMPPAPVPAGTPAPPGPATMMPDGTLGLTPPTTERFGQAATMEGIGAIGGTPPAFNRAAPGAE  462
cons             397 ***********:************************************************.****:  462

hamster_NONO     463 FAPNKRRRY  471
human_NONO       463 FAPNKRRRY  471
cons             463 *********  471
```

**Figure S2: Sequence alignment of hamster and human NONO.** Domains are coloured according the structure in Figure 1. In the consensus row identity is indicated as (*), low similarity as (.), and high similarity as (:).

```
hamster_SFPQ      1 MSRDRFRSRGGGGGGFHRRGGGGGRGGLHDFRSPPPGMGLNQNRGPMGPGPG--GPKPPIPPPPPH   64
human_SFPQ        1 MSRDRFRSRGGGGGGFHRRGGGGGRGGLHDFRSPPPGMGLNQNRGPMGPGPGQSGPKPPIPPPPPH   66
cons              1 ****************************************************  ************   66

hamster_SFPQ     65 QQQPQQPPPQQPPPQQPPPHQQPPPHQPPHQQ--PPPPPQDSSKPVVPQGPGSAPGVSPAPPPAGS  128
human_SFPQ       67 QQQ-QQPPPQQPPPQQPPPHQ-PPPHPQPHQQQQPPPPPQDSSKPVVAQGPGPAPGVGSAPPASSS  130
cons             67 *** ***************** ****  ****  *************.****.****..***.:.*  132

hamster_SFPQ    129 APPANPPTTGAPPG--PGPTPTPPPAVTSATPGPPPPSTPSSGVSTTPPQSGGPPPPPAGGAGPGP  192
human_SFPQ      131 APPATPPTSGAPPGSGPGPTPTPPPAVTSAPPGAPPPTPPSSGVPTTPPQAGGPPPPPAAVPGPGP  196
cons            133 ****.***:*****  **************.**.***:.*****.*****:********. .****  198

hamster_SFPQ    193 KQGPGPGPGGPKGGKMPGGPKPGGGPGMGAPGGHPKPPHRGGGEPRGGRQHHPPYHQQHHQGPPPG  258
human_SFPQ      197 GPKQGPGPGPGGPKGGKMPGGPKPGGGPGLSTPGGHPKPPHRGGGEPRGGRQHHPPYHQQHHQGPPPG  262
cons            199     **********************:.:*************************************  264

                                                                    RRM1
hamster_SFPQ    259 GPAARTEEKISDSEGFKANLSLLRRPGEKTYTQRCRLFVGNLPADITEDEFKRLFAKYGEPGEVFI  324
human_SFPQ      263 GPGGRSEEKISDSEGFKANLSLLRRPGEKTYTQRCRLFVGNLPADITEDEFKRLFAKYGEPGEVFI  328
cons            265 **..*:************************************************************  330

hamster_SFPQ    325 NKGKGFGFIKLESRALAEIAKAELDDTPMRGRQLRVRFATHAAALSVRNLSPYVSNELLEEAFSQF  390
human_SFPQ      329 NKGKGFGFIKLESRALAEIAKAELDDTPMRGRQLRVRFATHAAALSVRNLSPYVSNELLEEAFSQF  394
cons            331 ******************************************************************  396

                                         RRM2
hamster_SFPQ    391 GPIERAVVIVDDRGRSTGKGIVEFASKPAARKAFERCSEGVFLLTTTPRPVIVEPLEQLDDEDGLP  456
human_SFPQ      395 GPIERAVVIVDDRGRSTGKGIVEFASKPAARKAFERCSEGVFLLTTTPRPVIVEPLEQLDDEDGLP  460
cons            397 ******************************************************************  462

                                       NOPS
hamster_SFPQ    457 EKLAQKNPMYQKERETPPRFAQHGTFEYEYSQRWKSLDEMEKQQREQVEKNMKDAKDKLESEMEDA  522
human_SFPQ      461 EKLAQKNPMYQKERETPPRFAQHGTFEYEYSQRWKSLDEMEKQQREQVEKNMKDAKDKLESEMEDA  526
cons            463 ******************************************************************  528

                                                 coiled-coil domain
hamster_SFPQ    523 YHEHQANLLRQDLMRRQEELRRMEELHSQEMQKRKEMQLRQEEERRRREEEMMIRQREMEEQMRRQ  588
human_SFPQ      527 YHEHQANLLRQDLMRRQEELRRMEELHNQEMQKRKEMQLRQEEERRRREEEMMIRQREMEEQMRRQ  592
cons            529 ***************************.**************************************  594

hamster_SFPQ    589 REESYSRMGYMDPRERDMRMGGGGTMNMGDPYGSGGQKFPPLGGGGGIGYEANPGVPPATMSGSMM  654
human_SFPQ      593 REESYSRMGYMDPRERDMRMGGGGAMNMGDPYGSGGQKFPPLGGGGGIGYEANPGVPPATMSGSMM  658
cons            595 ************************:*****************************************  660

hamster_SFPQ    655 GSDMV-----------------RMIDVG---------------------  665
human_SFPQ      659 GSDMRTERFGQGGAGPVGGQGPRGMGPGTPAGYGRGREEYEGPNKKPRF  707
cons            661 ****                  * :. *                     709
```

**Figure S3: Sequence alignment of hamster and human SFPQ.** Domains are coloured according the structure in Figure 1. In the consensus row identity is indicated as (*), low similarity as (.), and high similarity as (:).

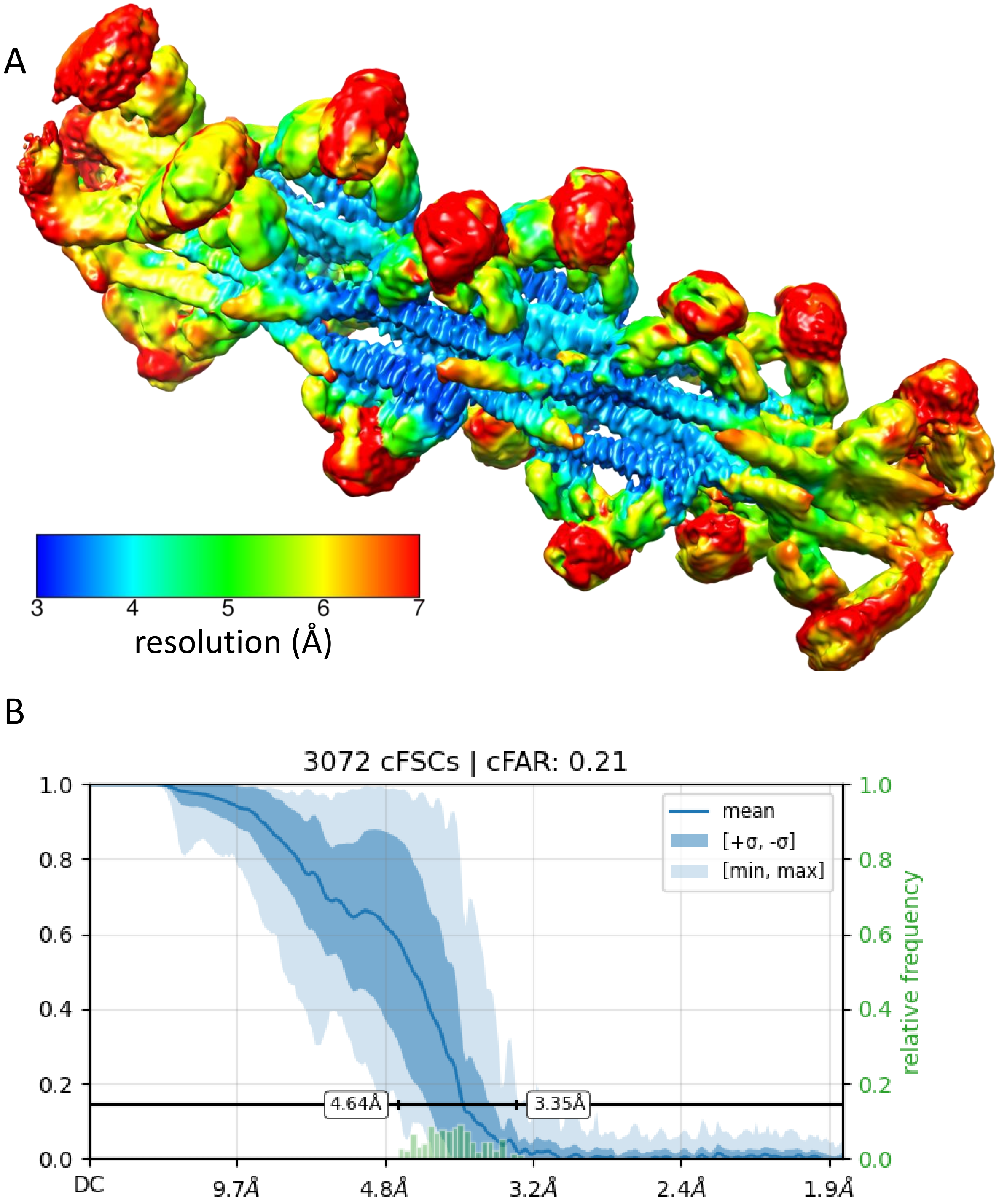


**Figure S4: Local and directional resolution of the filament.** (A) The map of the non-uniform refinement is coloured in the local resolution showing higher resolution for the coiled-coil, RRM2, and NOPS domains and low resolution for the RRM1 on the tip of the globular regions. (B) With the orientation diagnostics tool conical FSCs were analysed

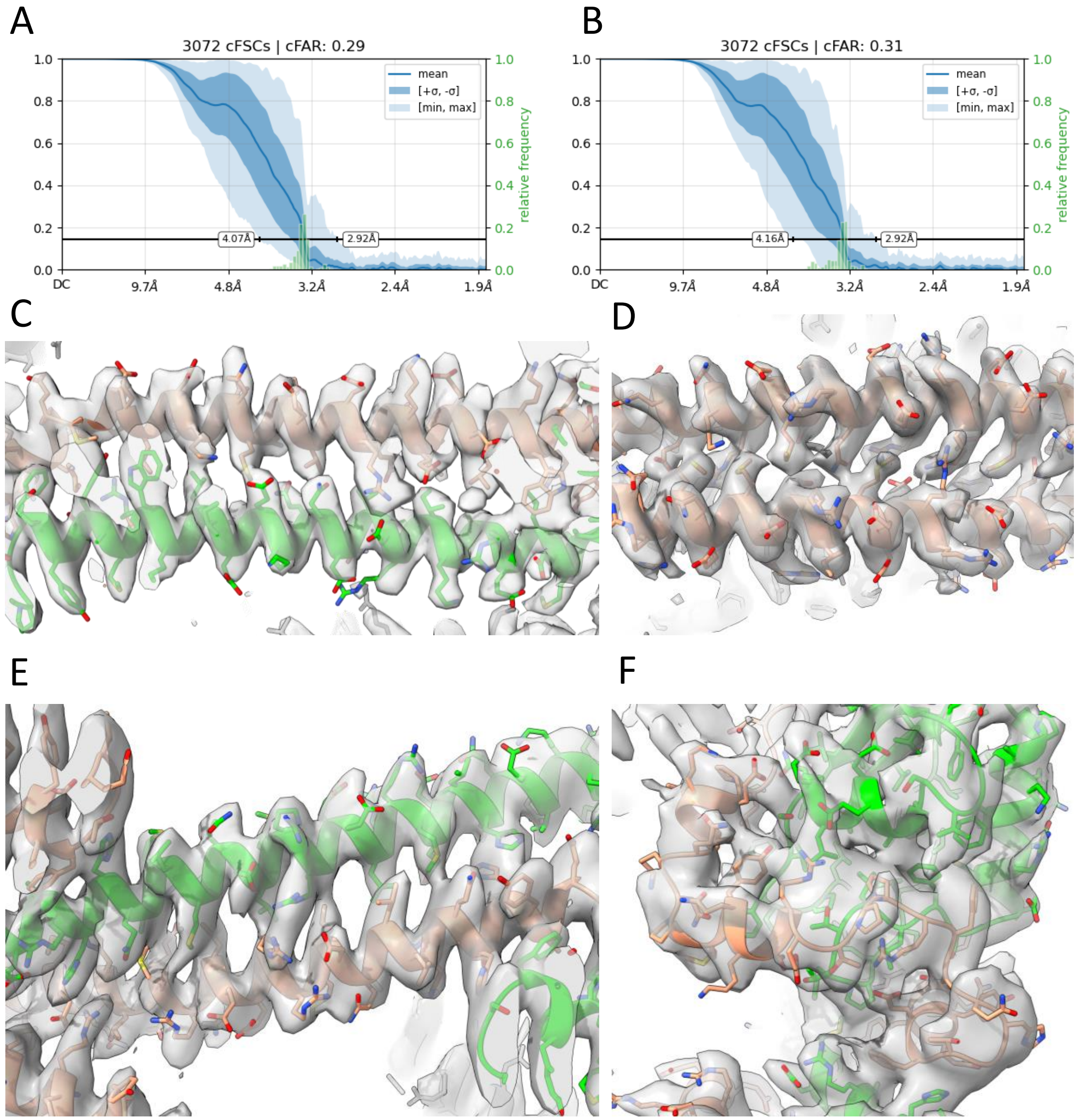


**Figure S5: Local refinements of the central units and modelling.** The directional resolutions are shown for local refinement of strand 1 (A) and strand 2 (B). Example densities and the model are shown for the local refinement of strand 2 where NONO is coloured green and SFPQ brown: (C) the dimerization domain of subunits AA/AB, (D) region 2 of subunits AA/CG, (E) region 1 of subunits AA/BB, and (F) RRM2 and NOPS of subunits BA/BB. The map is shown at a contour level of 10.

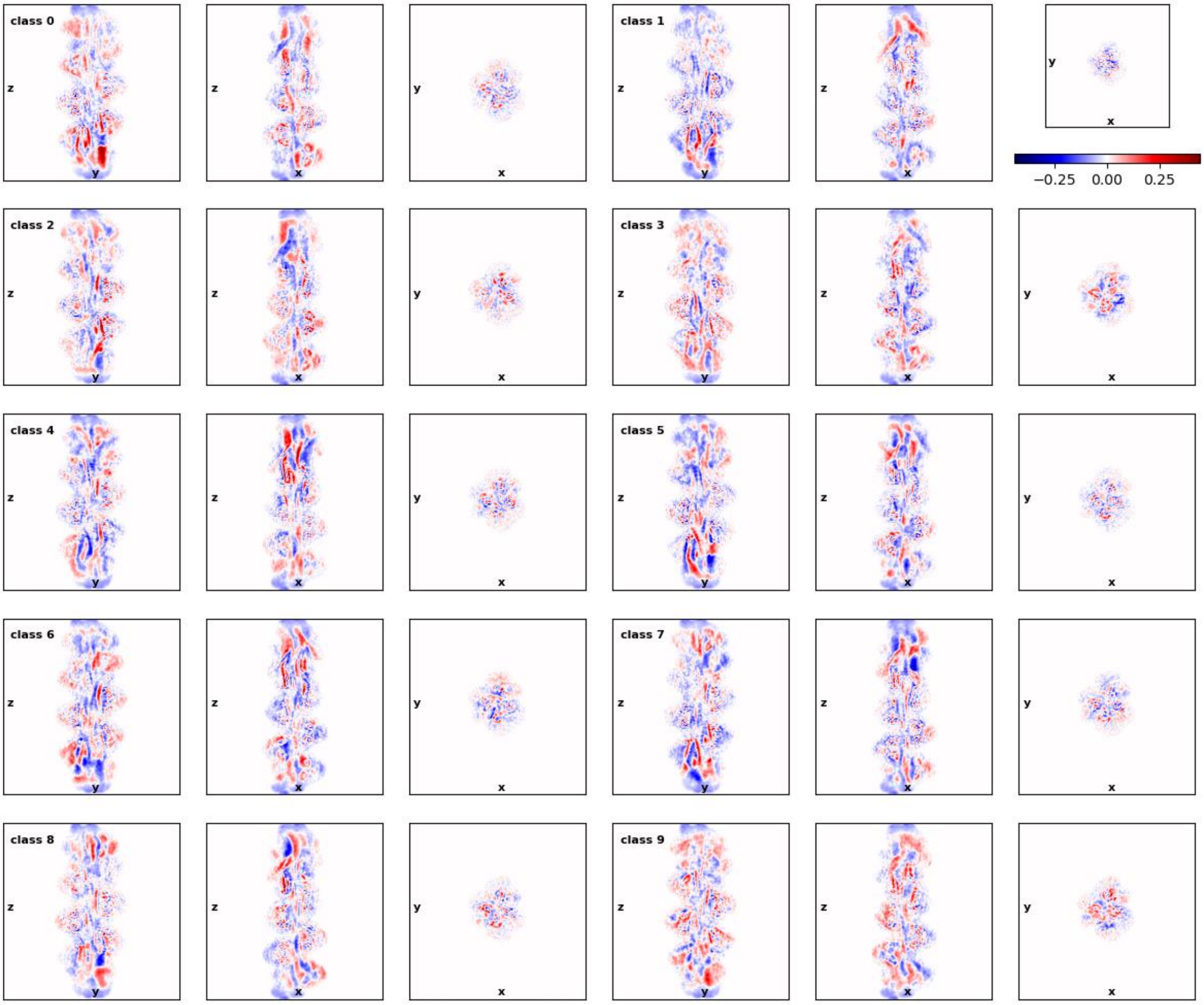


**Figure S6: 3D classification.** Real-space differences from the consensus for the 10 classes are shown. The classification was performed with CryoSparc 4.5.

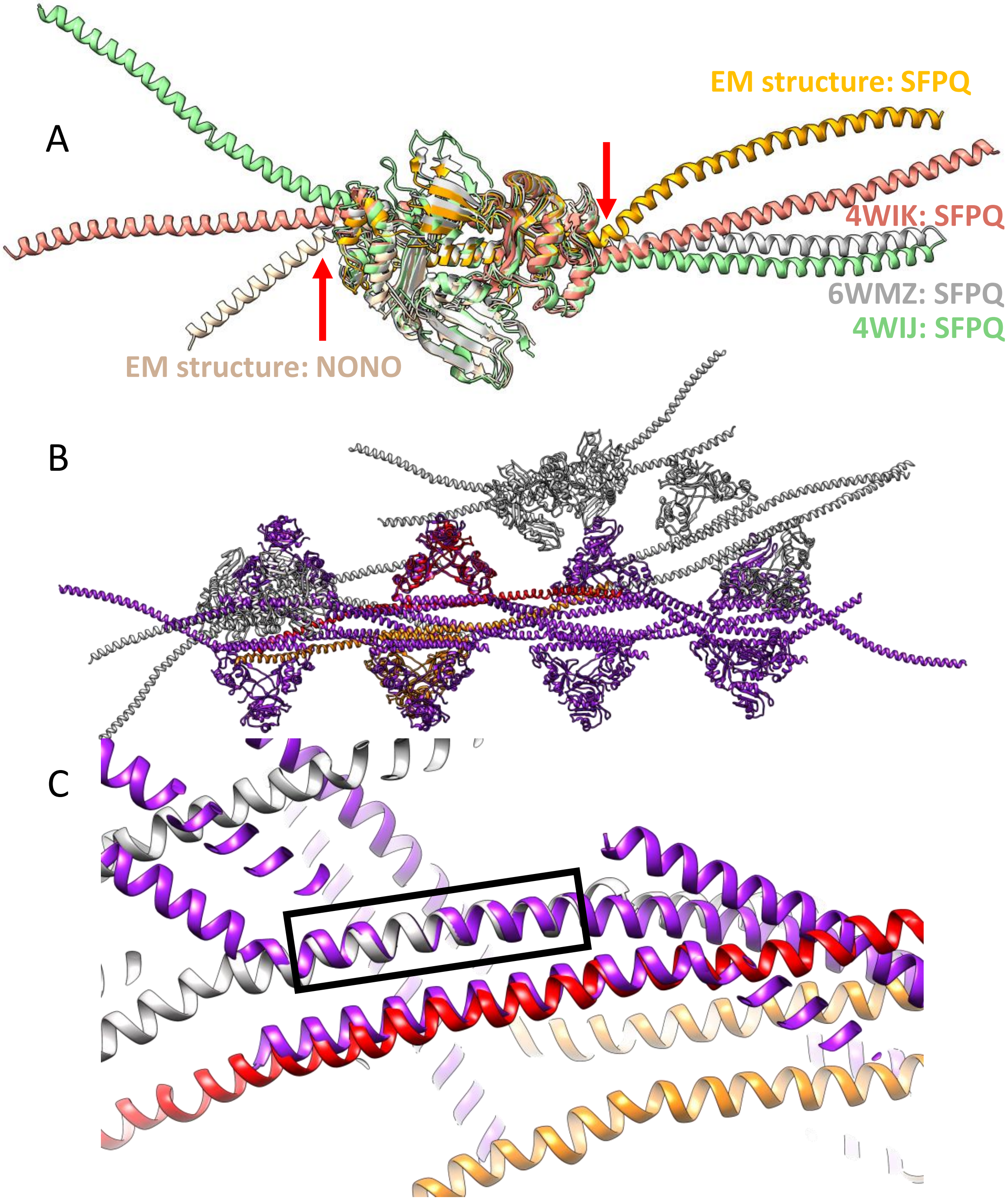


**Figure S7: Kink in the long α-helix:** (A) SFPQ shows a pronounced kink (red arrows) at S517 in the filamentous structure (orange) which is only small in the crystal structures of SFPQ (PDB: 4WIJ, 4WIK; green and red) and NONO/SFPQ (PDB: 6WMZ; blue). Also NONO displays a similar kink at residue E299 in the filamentous structure (light brown). (B) SFPQ homodimers in the unit cell of the crystal structure PDB 4WIJ (grey). One globular domain in the unit cell (red; until residue 516) was aligned to a dimer subunit of the filament (purple). Another dimer of the crystal structure (orange) has a similar offset and distance than dimers in the filament but the angle along the filament is different. (C) Region 1 of SFPQ (residues 523-545) of the filament (purple) where aligned with the crystal structure (grey, red, orange). The second helix in the coiled-coil is significant closer in the filament.

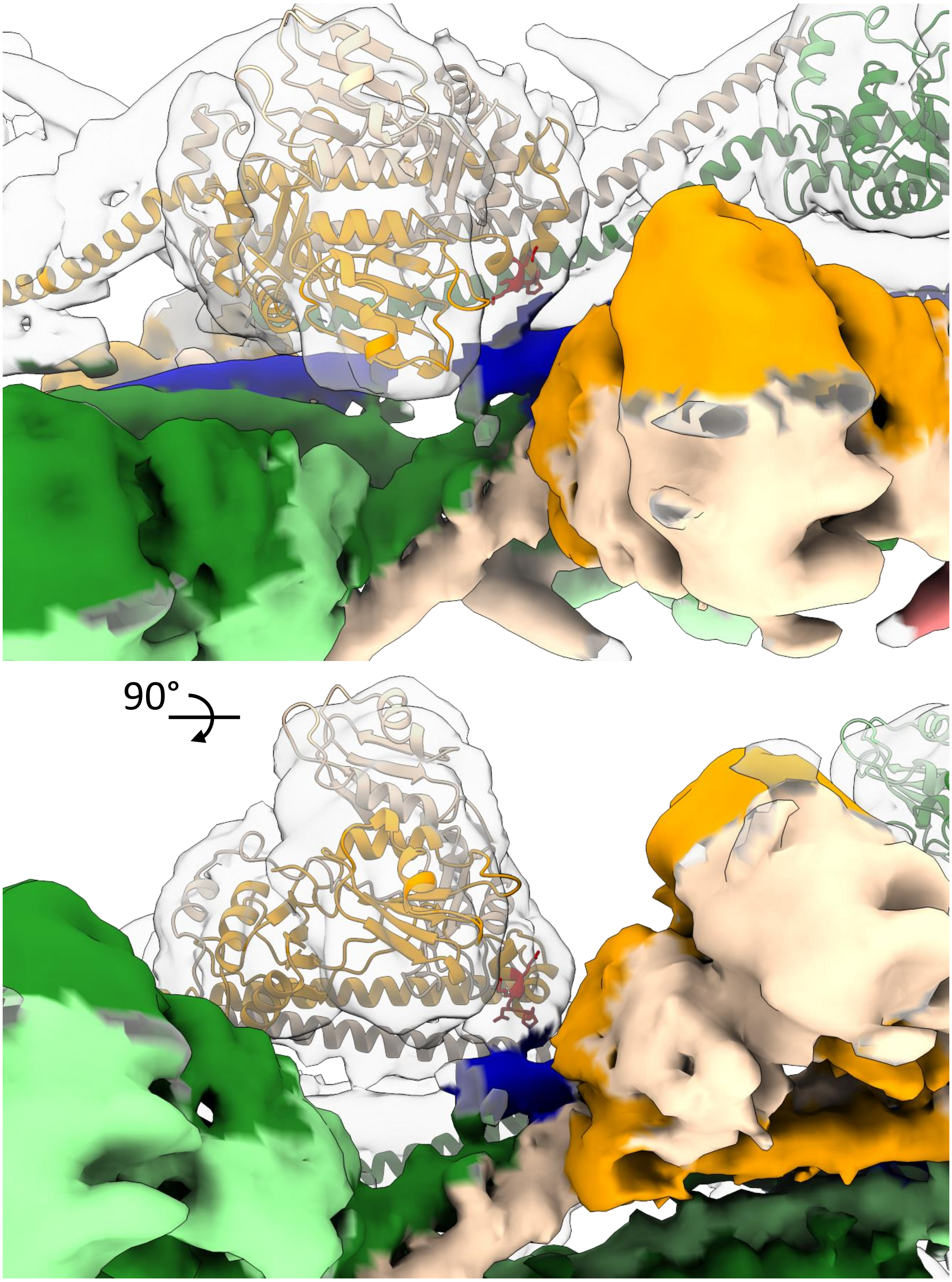


**Figure S8: Interstrand contact point.** The strand contact is shown here in a different way as in figure 4C. The NOPS domain of SFPQ (yellow model) at the loop Q465-P468 (red) is in contact to the end of the coiled-coil domain (solid blue) from a SFPQ of the other strand.

```
                             RRM1
hamster_NONO   1 MQSNKTFNLEKQNH------------------------------TP--------------------  16
hamster_SFPQ   1 MSRDRFRSRGGGGGGFHRRGGGGGRGGLHDFRSPPPGMGLNQNRGPMGPGPGGPKPPIPPPPPHQQ  66
cons           1 *. ::  .    .                                *                      66

hamster_NONO  17 ------------RKHHQHHHQQHHQQQQQQQQP---------------------------------  37
hamster_SFPQ  67 QPQQPPPQQPPPQQPPPHQQPPPHQPPHQQPPPPPQDSSKPVVPQGPGSAPGVSPAPPPAGSAPPA 132
cons          67             ::   *::   **  :**  *                                 132

hamster_NONO  38 -----------------------------------------------PPPIP-------------- 42
hamster_SFPQ 133 NPPTTGAPPGPGPTPTPPPAVTSATPGPPPPSTPSSGVSTTPPQSGGPPPPPAGGAGPGPKQGPGP 198
cons         133                                                *** *               198

hamster_NONO  43 ---------------------------------------------------------------ANG  45
hamster_SFPQ 199 GPGGPKGGKMPGGPKPGGGPGMGAPGGHPKPPHRGGGEPRGGRQHHPPYHQQHHQGPPPGGPAART 264
cons         199                                                                *.  264
                                                           RRM1
hamster_NONO  46 QQASSQNEGLTIDLKNFRKPGEKTFTQRSRLFVGNLPPDITEEEMRKLFEKYGKAGEVFIHKDKGF 111
hamster_SFPQ 265 EEKISDSEGFKANLSLLRRPGEKTYTQRCRLFVGNLPADITEDEFKRLFAKYGEPGEVFINKGKGF 330
cons         265 ::  *:.**:. :*. :*:*****:***.********.****:*:::** ***:.*****:*.*** 330

hamster_NONO 112 GFIRLETRTLAEIAKVELDNMPLRGKQLRVRFACHSASLTVRNLPQYVSNELLEEAFSVFGQVERA 177
hamster_SFPQ 331 GFIKLESRALAEIAKAELDDTPMRGRQLRVRFATHAAALSVRNLSPYVSNELLEEAFSQFGPIERA 396
cons         331 ***:**:*:******.***: *:**:******* *:*:*:****. ************ ** :*** 396
                                   RRM2
hamster_NONO 178 VVIVDDRGRPSGKGIVEFSGKPAARKALDRCSEGSFLLTTFPRPVTVEPMDQLDDEEGLPEKLVIK 243
hamster_SFPQ 397 VVIVDDRGRSTGKGIVEFASKPAARKAFERCSEGVFLLTTTPRPVIVEPLEQLDDEDGLPEKLAQK 462
cons         397 *********.:*******:.*******::***** ***** **** ***::*****:******. * 462
                      NOPS
hamster_NONO 244 NQQFHKEREQPPRFAQPGSFEYEYAMRWKALIEMEKQQQDQVDRNIKEAREKLEMEMEAARHEHQV 309
hamster_SFPQ 463 NPMYQKERETPPRFAQHGTFEYEYSQRWKSLDEMEKQQREQVEKNMKDAKDKLESEMEDAYHEHQA 528
cons         463 *  ::**** ****** *:*****: ***:* ******::**::*:*:*::*** *** * ****. 528
                                coiled-coil domain
hamster_NONO 310 MLMRQDLMRRQEELRRMEELHNQEVQKRKQLELRQEEERRRREEEMR---RQQEEMMRRQQ-EGF- 370
hamster_SFPQ 529 NLLRQDLMRRQEELRRMEELHSQEMQKRKEMQLRQEEERRRREEEMMIRQREMEEQMRRQREESYS 594
cons         529  *:******************.**:****:::**************    *: ** ****: *.:  594

hamster_NONO 371 KGTFPDAREQEIRMG---QMAMGGAMGINNRGAMPPAPVPTGTPAPPGPATMMPDGTLGLTPPTTE 433
hamster_SFPQ 595 RMGYMDPRERDMRMGGGGTMNMGDPYGSGGQ-KFPPLGGGG---------GIGYEANPGVPPATMS 650
cons         595 :  : *.**:::***    * **.. * ..:  :**               :  :.. *:.*.* . 660

hamster_NONO 434 RFGQAATMEGIGAIGGTPPAFNRPAPGADFAPNKRRRY 471
hamster_SFPQ 651 GSMMGSDMVRMIDV-----------------------G 665
cons         661     .: *  :  :                         698
```

**Figure S9: Sequence alignment of hamster NONO and SFPQ.** Domains are coloured according the structure in figure 1. In the consensus row identity is indicated as (*), low similarity as (.), and high similarity as (:). The histidine rich motif in NONO is marked in yellow.

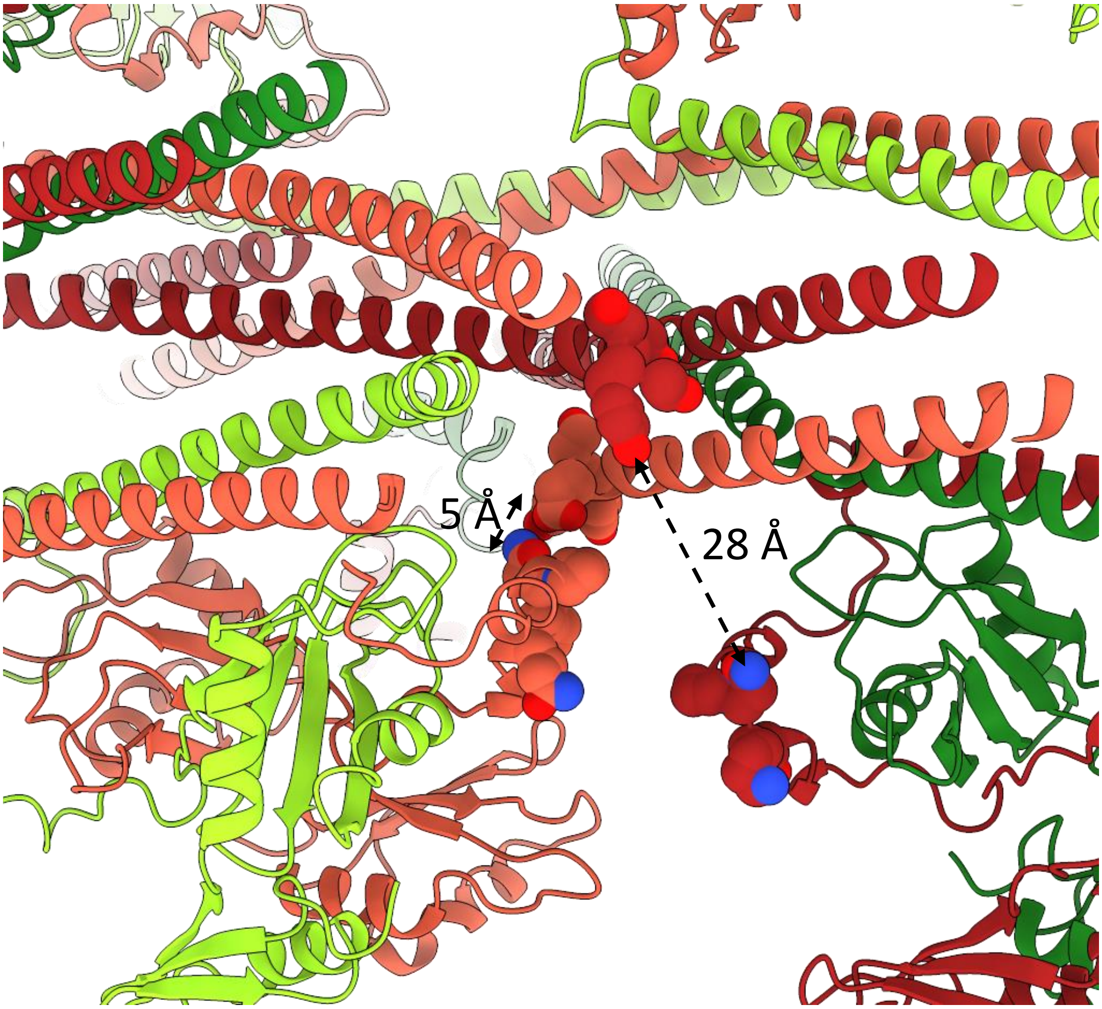


**Figure S10: Interstrand contact point.** The end of the long α-helix of SFPQ (R591-A594) is shown as spheres as well as the contact loop of the NOPS domain (Q461-P464). For the light red SFPQ subunits the interstrand contact is established but not for the dark red SFPQ pair. Here the contact is blocked by the long helix of the light red SFPQ. Coloring as in Figure S4.

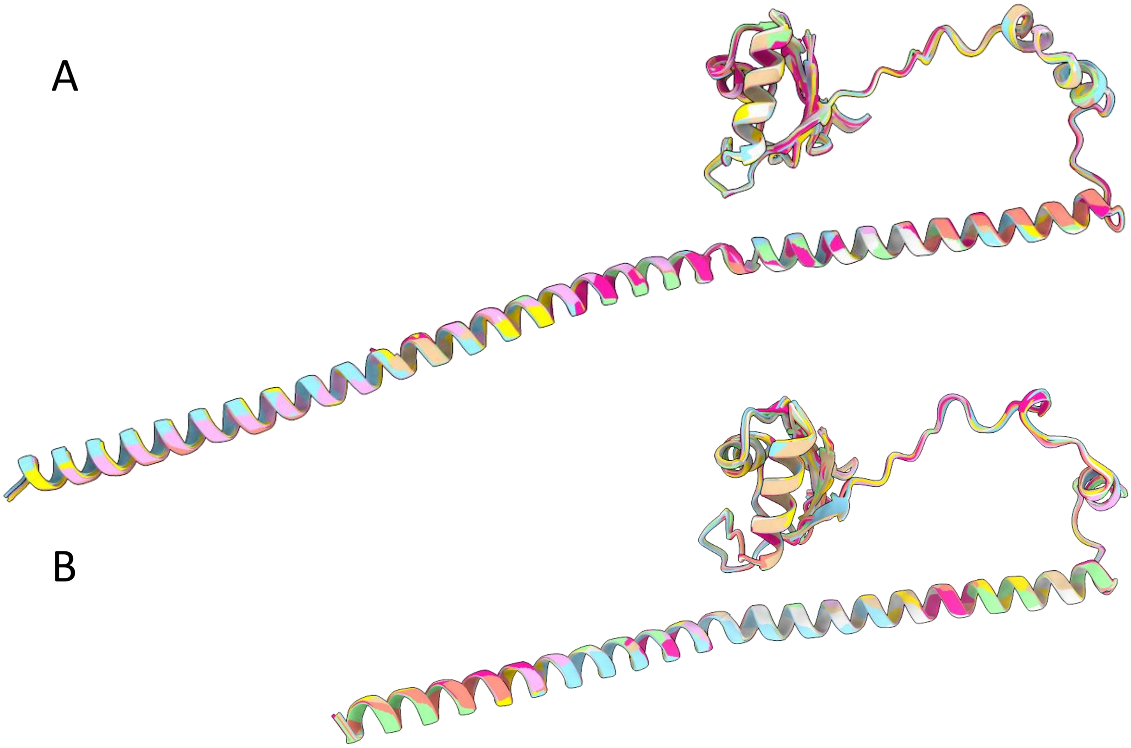


**Figure S11: Overlay of subunits from the local refinements.** Using ChimeraX matchmaker, subunits of SFPQ (A) or NONO (B) were overlayed (PDB 9GLC and 9GLD). No significant differences in the backbone can be seen although SFPQ subunits have two different environments.
